# Combination therapy with nirmatrelvir and molnupiravir improves the survival of SARS-CoV-2 infected mice

**DOI:** 10.1101/2022.06.27.497875

**Authors:** Ju Hwan Jeong, Santosh Chokkakula, Seong Cheol Min, Beom Kyu Kim, Won-Suk Choi, Sol Oh, Yu Soo Yun, Da Hyeon Kang, Ok-Jun Lee, Eung-Gook Kim, Jang-Hoon Choi, Joo-Yeon Lee, Young Ki Choi, Yun Hee Baek, Min-Suk Song

## Abstract

As the SARS-CoV-2 pandemic remains uncontrolled owing to the continuous emergence of variants of concern, there is an immediate need to implement the most effective antiviral treatment strategies, especially for risk groups. Here, we evaluated the therapeutic potency of nirmatrelvir, remdesivir, and molnupiravir and their combinations in SARS-CoV-2-infected K18-hACE2 transgenic mice. Systemic treatment of mice with each drug (20 mg/kg) resulted in slightly enhanced antiviral efficacy and yielded an increased life expectancy of only about 20–40% survival. However, combination therapy with nirmatrelvir (20 mg/kg) and molnupiravir (20 mg/kg) in lethally infected mice showed profound inhibition of SARS-CoV-2 replication in both the lung and brain and synergistically improved survival times up to 80% compared to those with nirmatrelvir (P= 0.0001) and molnupiravir (P= 0.0001) administered alone. This combination therapy effectively reduced clinical severity score, virus-induced tissue damage, and viral distribution compared to those in animals treated with these monotherapies. Furthermore, all these assessments associated with this combination were also significantly higher than that of mice receiving remdesivir monotherapy (P= 0.0001) and the nirmatrelvir (20 mg/kg) and remdesivir (20 mg/kg) combination (P= 0.0001), underscored the clinical significance of this combination. By contrast, the nirmatrelvir and remdesivir combination showed less antiviral efficacy, with lower survival compared to nirmatrelvir monotherapy, demonstrating the inefficient therapeutic effect of this combination. The combination therapy with nirmatrelvir and molnupiravir contributes to alleviated morbidity and mortality, which can serve as a basis for the design of clinical studies of this combination in the treatment of COVID-19 patients.

**IMPORTANCE:** Since SARS-CoV-2 spread rapidly with the emergence of new variants of concerns, it is necessary to develop effective treatment strategies to treat elderly individuals and those with comorbidities. Antiviral therapy using a combination of drugs is more effective in eradicating viruses and will undoubtedly improve the clinical outcome and survival probability of hospitalized SARS-CoV-2 patients. In the current study, we observed three FDA-approved antivirals nirmatrelvir, remdesivir, and molnupiravir have therapeutic significance with moderate survival for their monotherapies against SARS-CoV-2 infected K18-hACE2 mouse model. The combination of nirmatrelvir and molnupiravir showed significant antiviral activity and a higher survival rate of approximately 80%, providing *in vivo* evidence of the potential utility of this combination. In contrast, nirmatrelvir and remdesivir combination showed less antiviral potency and emphasized the ineffective significance with less survival. The current study suggests that the nirmatrelvir and molnupiravir combination is an effective drug regimen strategy in treating SARS-CoV-2 patients.

## INTRODUCTION

Since severe acute respiratory syndrome coronavirus-2 (SARS-CoV-2) was first identified in China in December 2019, the virus has spread rapidly worldwide, causing 538,321,874 confirmed cases, including 6,320,599 deaths (1). It poses a severe threat to health care systems and economies and is devastating to some populations, such as elderly individuals and those with comorbidities. We can also expect to coexist with the virus for a long time. Fortunately, well-established technology platforms and unprecedented efforts have led to the development of an effective vaccine as fast as the disease spread, saving the lives of millions and billions of people. The vaccine is also effective for multiple variants, such as alpha, beta, gamma, delta, and omicron, that have emerged thus far (2).

Despite the availability of vaccines, it is imperative to develop effective drugs that have broad-spectrum effects on current and future potential coronavirus infections, which could provide a way to ease disease symptoms and prevent death. Several direct-acting antiviral agents have been approved by regulatory agencies and are advancing in various stages of clinical development. These antivirals target different viral proteins and are categorized into three classes. Monoclonal antibodies (mAbs) are prescribed as drugs that target the spike protein, and direct-acting small molecules interfere with the viral replication machinery. Usually, mAbs target only the spike protein, the timeline for approval of a novel antibody for viral infection management is prolonged, and side effects such as antibody-dependent enhancement of viral infections are also considered. Moreover, most of the approved mAbs are not sufficiently effective against SARS-CoV-2 variants (3, 4). Direct-acting small molecules can be divided into two classes: those targeting RNA-dependent RNA polymerase (RdRp) and those targeting viral proteases, such as the main protease (M^pro^, also known as 3CL^pro)^ and papain-like protease (PL^pro^). Remdesivir (RDV) is the prodrug of the nucleoside analog GS-441524, the first approved drug that targets RdRp (5, 6). Molnupiravir (MK-4482 or EIDD-2801), a prodrug of EIDD-1931 (β-D-N4-deoxycytidine), also targets RdRp and exhibits improved inhibition of several RNA viruses, including SARS-CoV-2 (7-9). M^pro^ is a cysteine protease that is a promising target. It is responsible for the catalytic cleavage of the conserved regions of polyproteins PP1a and PP1ab, and its blockade inhibits the synthesis of many nonstructural proteins that are crucial for viral proliferation (10). Nirmatrelvir and ritonavir are co-administered oral antiviral drugs with the trade name Paxlovid. This drug acts as an irreversible inhibitor of the M^pro^ of SARS-CoV-2.

Combination therapy with two distinct drugs has led to different results, such as antagonistic, synergistic, and additive effects (11). In general, the synergistic and additive effects of combination therapy are significant and are facilitated by targeting multiple pathways of the virus replication machinery. Viral diseases caused by human immunodeficiency virus (HIV), hepatitis C virus (HCV), and influenza are classic examples, for which combination therapy has become the benchmark treatment (12-16). Although combination therapy does not always end in a cure, it can significantly improve quality of life and prolong life because of increased therapeutic efficacy and prevention of the development of drug resistance. In addition to historical experience in the treatment of viral diseases, there have been some recent studies on combination therapy synergistically reducing the SARS-CoV-2 load (11, 17). Combination therapy has been suggested as potent and effective when it targets different mechanisms and processes of the virus life cycle, such as the proteases (M^pro^) and polymerases (RdRp) of SARS-CoV-2. Furthermore, although there are several ongoing clinical studies with monotherapies, very few clinical studies have used combination therapies. Moreover, *in vivo* information in the context of SARS-CoV-2 infection is essential before considering combination therapy for clinical studies. Small-animal models that accurately mimic human disease usually afford fundamental insights into the pathogenesis of SARS-CoV-2 infection (18-20). K18-hACE2 transgenic mice with the human angiotensin I-converting enzyme 2 (ACE2) receptor predominantly express hACE2 under the control of the cytokeratin-18 promoter (KRT18) and have been proposed as a model for the study of antiviral agents (21). In the context of hACE2 expression, mice have been shown to develop severe lung pathology and impaired lung function, with accompanying neuronal death, upon SARS-CoV-2 infection (18, 22).

Screening FDA-approved antiviral drugs for efficacy against SARS-CoV-2 will help reduce the global impact of the current pandemic. Additionally, such an approach strongly emphasizes the practical and clinical need to develop appropriate combination therapies and increase access to COVID-19 patients, especially for high-risk groups. Therefore, we performed an *in vivo* study using a SARS-CoV-2-infected hACE2 transgenic mouse model and evaluated the potential efficacy of the drugs nirmatrelvir (NTV), remdesivir (RDV), and molnupiravir (MPV) and their combinations, nirmatrelvir and remdesivir (NTV-RDV) and nirmatrelvir and molnupiravir (NTV-MPV). We summarized our therapeutic insights into these drug combinations, which are of clinical significance, and we provided an alternative prescription for a more effective drug regimen for treating SARS-CoV-2 patients.

## RESULTS

### Therapeutic efficacy of nirmatrelvir, remdesivir, and molnupiravir in SARS-CoV-2-infected mice

The treatment dose and infectious virus titer optimization in mice was defined by the criteria of the lowest dose with maximum therapeutic efficacy at the assigned time points. The suboptimal mouse lethal dose (MLD50) for the infected mice was determined by infecting the mice with various titers of Beta-CoV/Korea/KCDC03/2020 (Fig S1). We conducted two dose-optimization studies (10 and 20 mg/kg body weight) to determine the low-dose therapeutic efficacy using hACE-2 transgenic mice (Fig S1A and S1B). Based on the weight change, survival, viral load reduction, and pathology-related aspects, we determined that a 20 mg/kg body weight dose was optimal for low-dose therapeutic efficacy *in vivo* (Fig S2).

To evaluate the therapeutic efficacy of NTV, RDV, and MPV, mice were infected with 5 MLD50 of SARS-CoV-2 followed by administration via oral gavage of 20 mg/kg body weight NTV and MPV and intraperitoneal administration of RDV alone. Six hours after the onset of infection, treatment was started on the same day and continued twice daily for five consecutive days (Fig 1A). As body weight loss is a crude marker for viral pathogenesis, mice were monitored daily from day 0 (0 DPI) for body weight changes, and all mice continued to lose weight from 5 DPI and showed a maximum weight loss by 7 or 8 DPI. No protection against weight loss was observed in RDV-treated mice (20.5%) compared to that in mock-treated mice (21.5%) by 7 DPI, whereas approximately 5-6% protection against weight loss was observed in NTV-and MPV-treated animals (Fig 1B). Congruent with protection against weight loss, NTV (P= 0.0001) and MPV (P= 0.0001) produced significantly higher (36% and 43%) survival, respectively, while RDV (P=0.1172) produced a lower (21%) survival rate than that observed with mock treatment. The survival rates of the NTV-(P=0.0197) and MPV-treated (P=0.0055) animals were significantly higher than that of the RDV-treated animals (Fig 1C). In addition to the improvement in body weight and survival, the NTV-(P= 0.0001) and MPV-treated mice (P= 0.0001) exhibited significantly higher clinical scores than RDV-treated mice from 5 DPI (Fig 1D). We analyzed the therapeutic effect of these antivirals on the survival probability of mice. As changes in brain tissues and neurological involvement potentially contributed to the lethality of the K18-hACE2 mice, we focused on the brain along with the lung. Therefore, tissues were collected from both the lung and brain at 5 DPI for viral titer assessment and histopathological and immunohistochemical analysis. We assessed the replication capability of the virus in the brain and lungs via TCID50 and RT–qPCR. Titers reduced by 0.7, 0.4, and 1 log10 TCID50 were observed in the lung tissue of NTV (P=0.1987), RDV-(P=0.7062) and MPV-treated (P=0.05714) mice, respectively, compared to those in mock-treated mice (Fig 2A). We further evaluated the viral titer reduction among these treated animals and observed a better log10TCID50 titer reduction in the MPV-treated animals than in the NTV and RDV-treated animals (Fig 2A). The same scenario regarding the viral titer was also observed in brain tissue, which has low viral replication potential (Fig 2A). To determine the relationship between the viral titer and viral RNA copy number in tissue samples, we further performed RT–qPCR on parallel tissues used for TCID50. As with the TCID50, the same trend was observed in RNA copy numbers, where, compared to RDV-treated animals, MPV-treated animals showed significantly higher copy number reduction in the lung and brain; however, this reduction was not significantly higher compared to that observed in NTV-treated animals (Fig 2B and 2C). We evaluated the viral antigen levels by immunochemistry and histopathological changes determined by H&E staining of lung and brain tissues at 5 DPI. No potent decrease in the expression levels of the SARS-CoV-2 N protein was observed in the RDV-treated animals, whereas a reduction in viral distribution was observed in the NTV and MPV-treated animals, with a greater reduction in the MPV-treated animals (Fig 3A and 3B). H&E staining revealed reduced pulmonary damage in the alveolar and peribronchiolar regions of the lung slices of MPV-treated animals at 5 DPI (Fig 3A). Severe inflammation was observed in the brains of the mock-treated mice at 5 DPI, whereas, compared to the animals in the other groups, the NTV and MPV-treated animals exhibited alleviation of such changes (Fig 3B). All these findings demonstrate that MPV, a nucleotide analog that targets the conserved RdRp, is efficient in diminishing the infectious SARS-CoV-2 burden and is further associated with better survival compared with that with other monotherapy treatments.

**Figure 1:**
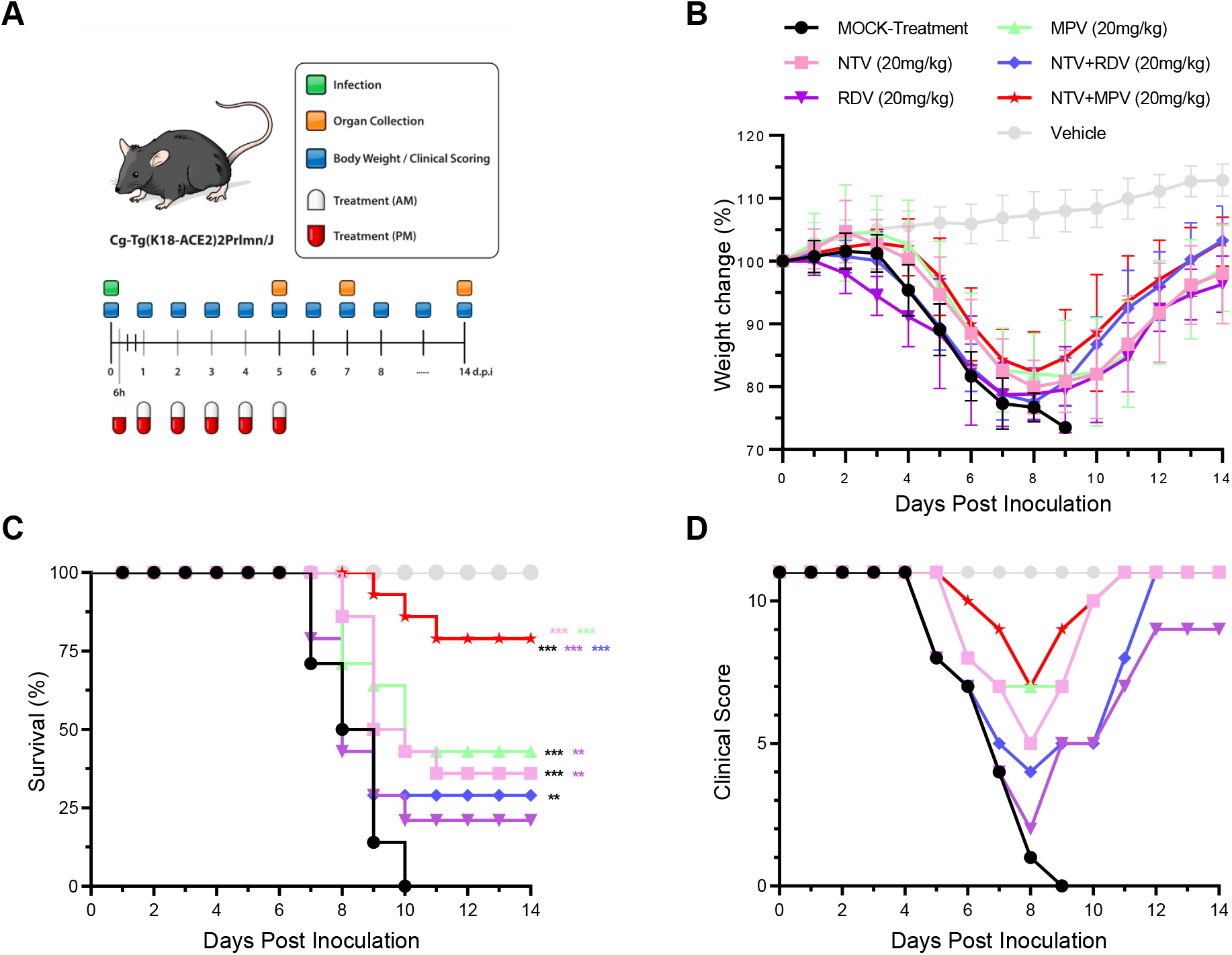
Evaluation of the therapeutic efficacy of nirmatrelvir, remdesivir, and molnupiravir monotherapies and their combinations, nirmatrelvir-remdesivir, and nirmatrelvir-molnupiravir, in SARS-CoV-2-infected K18-hACE2 transgenic mice. **(A)** Schematic illustration of the study design. Mice were infected with 5 MLD50 of SARS-CoV-2 followed by administration of 20 mg/kg drugs for 5 consecutive days. Disease progression and other clinical parameters were monitored until 14 DPI, and the brain and lung were collected at 5 DPI for viral titer estimation and histopathological and immunohistochemical analyses. **(B)** Body weight was monitored daily for 14 days and is expressed as a percentage relative to the initial body weight on day 0. **(C)** Kaplan–Meier plot of the survival of all the groups (log-rank Mantel-Cox; **p < 0.01, ***p < 0.001). **(D)** The clinical score was evaluated by assessing appearance (fur, eye closure), activity, and movement from 0 DPI to 14 DPI, and the cumulative clinical score of mice in each group is indicated.

**Figure 2:**
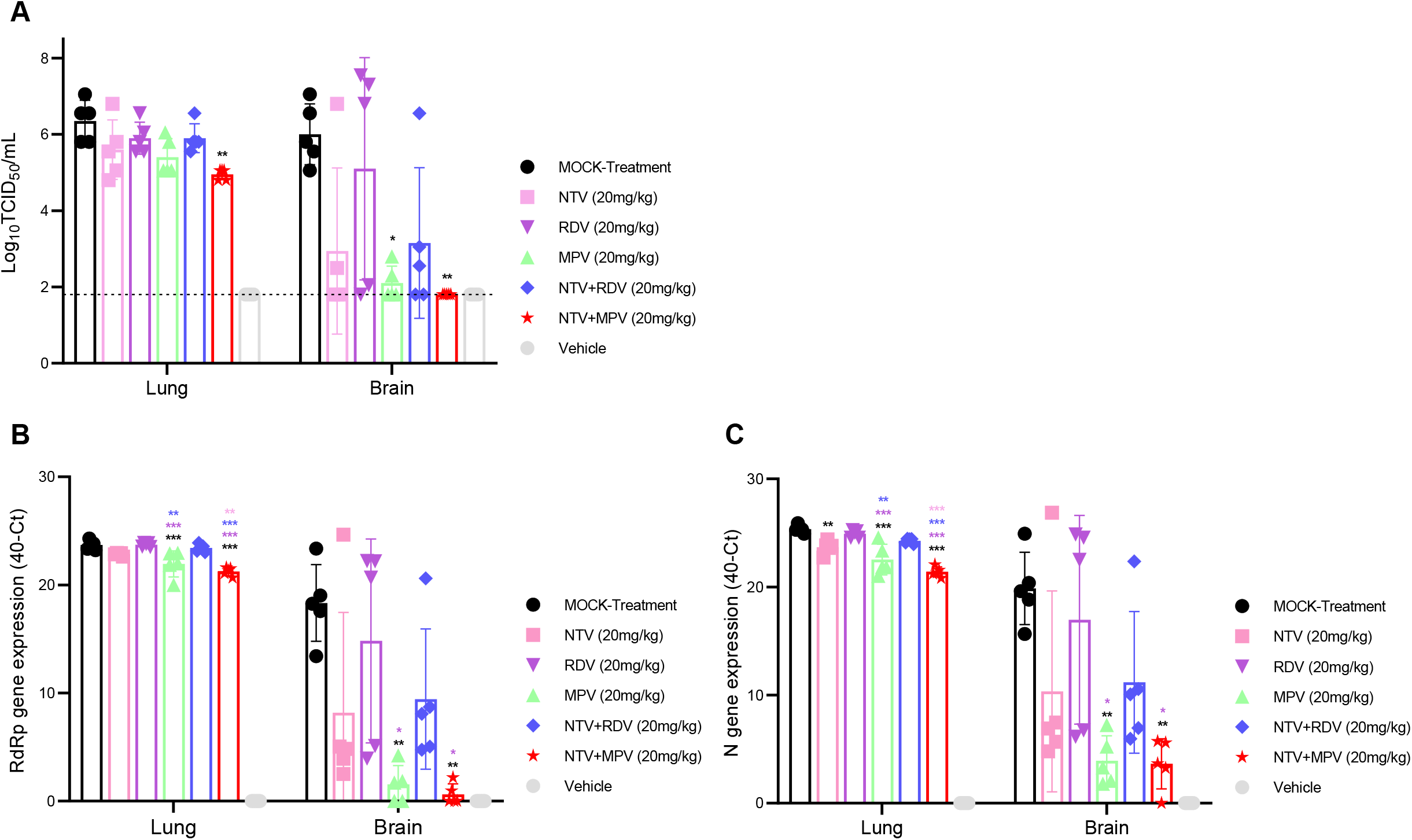
Evaluation of the viral load in SARS-CoV-2-infected mice treated with nirmatrelvir, remdesivir, and molnupiravir monotherapies, their combinations, nirmatrelvir-remdesivir, and nirmatrelvir-molnupiravir. **(A)** Lungs and brains were collected at 5 DPI, and viral titers were estimated. The viral titer is expressed in log10 PFU/mL, and the limit of detection is shown in a dotted line (one-way ANOVA with Tukey’s multiple comparison test; *p < 0.05**p < 0.01). **(B)** Viral RdRp gene quantification in both the lungs and brain at 5 DPI was determined via RT–qPCR and results are expressed in 40-Ct (One-way ANOVA with Tukey’s multiple comparison test; *p < 0.05, **p < 0.01, ***p < 0.001). **(C)** Viral N gene quantification in both the lung and brain at 5 DPI via RT–qPCR. The data are expressed in 40-Ct (one-way ANOVA with Tukey’s multiple comparison test; *p < 0.05, **p < 0.01, ***p < 0.001).

**Figure 3:**
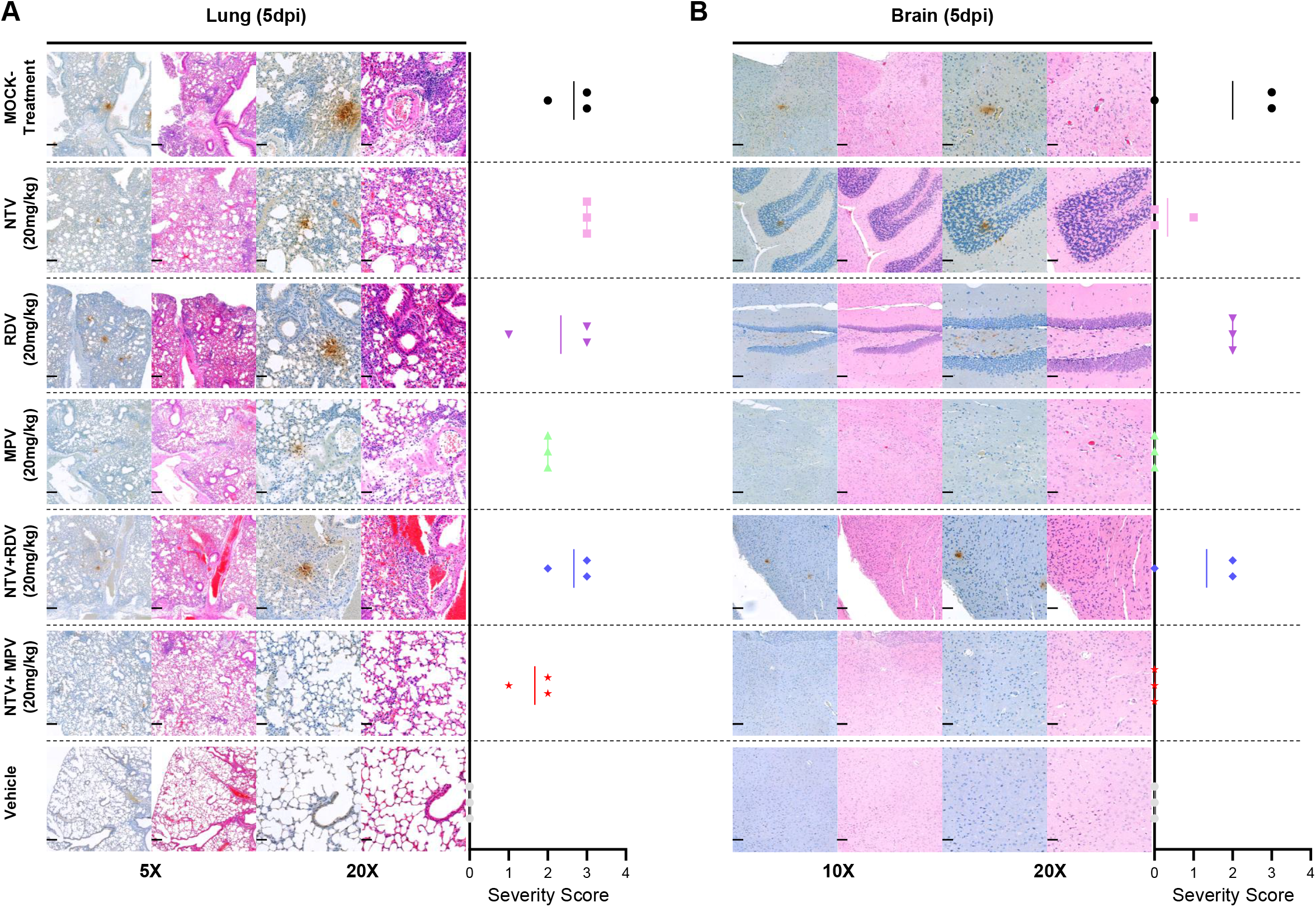
Histopathology and immunohistochemistry analysis of SARS-CoV-2-infected mice treated with 20 mg/kg body weight nirmatrelvir, remdesivir, and molnupiravir monotherapy and their combinations, nirmatrelvir-remdesivir, and nirmatrelvir-molnupiravir. **(A)** Pathological changes and virus distribution were measured at 5 DPI in the lung tissue by H&E staining and immunohistochemical staining with an anti-nucleocapsid antibody, respectively. Images are shown at both low (5X) and high (20X) power resolution. **(B)** Neuronal death and virus distribution were measured at 5 DPI in brain tissue by H&E staining and immunohistochemistry, respectively. Images are shown at low (10X) and high (20X) power resolutions.

### The combination of nirmatrelvir and remdesivir demonstrated less antiviral activity, with a lower survival rate, in an in vivo model

Combination treatment is considered more effective than single-drug treatment, so we examined the therapeutic benefits of NTV and RDV combination therapy in SARS-CoV-2-infected mice. Unfortunately, the combination of NTV and RDV did not result in any significant benefits and yielded the same or less antiviral action as NTV monotherapy. Despite the lack of superior protection afforded by this combination regarding weight loss compared to that with a mock-treatment and RDV treatment (Fig 1B), it resulted in approximately 7% lower survival compared to that with NTV monotherapy (Fig 1C). As shown in Figure 2, it also failed to inhibit viral replication in both the lungs and the brain and thus produced antagonistic activity compared to that of NTV monotherapy (Fig 2). Although the overall pathological changes in the lungs and brain of the combination-treated mice were similar to those of mock-treated and RDV-treated mice at 5 DPI, the combination produced an inefficient therapeutic effect (Fig 3).

### Nirmatrelvir and molnupiravir combination therapy conferred enhanced survival to SARS-CoV-2-infected mice

Since only modest survival benefits were observed with the NTV-RDV combination, we further tested another combination with the same therapeutic targets, NTV and MPV, to determine if the result would be functionally synergistic, additive, or optimal. Hence, we administered NTV together with MPV (20 mg/kg body weight each) to mice and observed more promising results than those with either of the drugs administered alone. Along with the NTV and MPV single-drug treatments, the NTV-MPV combination treatment also exhibited protective efficacy against weight loss (15.5%) compared to that mock-treatment (21.5%) (P=0.0176) at 7 DPI (Fig 1B). This protection against weight loss was 4.7-5.5% better than that with RDV and the NTV-RDV combination; however, this effect was neither synergistic nor significant in animals that received the NTV or MPV drugs alone (Fig 1B). Furthermore, in addition to enhanced protection against body weight loss, mice treated with this combination showed a rapid improvement in body weight recovery compared to that of animals treated with other drugs. Remarkably, the NTV-MPV combination drug-treated animals did not exhibit any rapid decline in the clinical score, as did other treated animals (Fig 1D). Additionally, a rapid increase in the clinical score was observed from 8 DPI in the NTV-MPV combination group, while other groups maintained only moderate progress in their clinical score (Fig 1D). The optimal survival probability was exhibited by NTV, RDV, MPV, and NTV-RDV-treated animals, which was largely improved to more than 80% with NTV-MPV combination therapy, exhibiting synergetic and significant protection compared to that with NTV (P=0.0001) or MPV (P=0.0001) monotherapy (Fig 1C). Similarly, the improvement in survival associated with this combination was substantially greater than that associated with RDV monotherapy (P= 0.0001) or NTV-RDV combination (P= 0.0001) treatment. When we compared the viral replication with this combination in the lungs and brain with that with mock and other treatment groups, we found that the combination significantly reduced viral replication. The NTV-MPV combination reduced the log10 TCID50 viral titers by 1.4 at 5 DPI in the lung (P=0.0019), whereas the log10 TCID50 viral titer was reduced by 4.2 in the brain (P=0.0091) compared to that with mock treatment (Fig 2A). Additionally, with the combination therapy, the virus titer was reduced by 0.65, 0.9, 0.45, or 0.95 log10 TCID50 in the lung compared to that with the NTV, RDV, or MPV monotherapy or NTV-RDV combination, respectively. We further observed significant viral inhibition by this combination in the brain tissue compared with that with the mock-treatment; however, the difference between NTV and MPV monotherapy was not significant. The viral RNA expression was significantly inhibited in the NTV-MPV-treated group than in the other treatment groups except for MPV-treated for both RdRp and N genes in the lung, whereas it was only significant with the mock and RDV-treated groups in the brain (Fig 2B and 2C). Excessive viral titer reduction in the brains of NTV-MPV-treated mice compared to other groups revealed that administration of the NTV-MPV combination may be more beneficial than other therapies in reducing the lung virus load and preventing the spread of SARS-CoV-2 beyond the respiratory system. The NTV-MPV combination-treated groups exhibited greater recovery of lung impairment, lung inflammation, and other related issues at 5 DPI than the other treated groups (Fig 3A). None to moderate reduction of bronchiolitis, alveolitis, and encephalitis with neutrophils and macrophages was observed in the monotherapy and NTV-RDV treated groups, whereas effective reduction of all these lesions was observed in the NTV-MPV combination-treated animals at 5 DPI (Fig 3).

## DISCUSSION

Despite the widespread availability of vaccines, the unrestrained SARS-CoV-2 pandemic needs advancement of effective antiviral therapy development to adeptly target most of the emerging SARS-CoV-2 strains and further support clinical outcomes with greater clinical utility, especially for high-risk groups (23). In October 2020, the FDA approved RDV and recommended the treatment of hospitalized patients with severe COVID-19 in many countries. These recommendations are partly based on the belief that the use of RDV could reduce recovery time, allowing faster discharge of patients from hospitals during the pandemic. In contrast, several clinical studies have highlighted the lack of survival benefits associated with RDV and the WHO has recently recommended against the use of RDV for hospitalized patients. Later, MPV was recognized for its therapeutic and prophylactic potential and was recently approved for emergency use by regulatory agencies. Significant clinical benefits have been observed for patients who have been administered MPV orally (8, 24-26). NTV is an essential component of Paxlovid, which impedes the main protease of SARS-CoV-2 and has recently been reported to reduce the risk of hospitalization or death in COVID-19 patients by 89% (27, 28). Antiviral therapy with a combination of drugs that can target viral replication and entry machinery components is more efficient in eradicating viruses and can undoubtedly improve the clinical outcomes and survival probability of hospitalized patients. As such, we evaluated the therapeutic efficacy of NTV, RDV, and MPV and their combinations in SARS-CoV-2-infected K18-hACE2 transgenic mice and demonstrated the superior antiviral activity of the NTV-MPV combination compared with those of NTV, MPV, RDV, and NTV-RDV.

Antivirals such as RDV, MPV, and Paxlovid target highly conserved enzymes involved in virus replication that have remained relatively invariant as new VOCs have emerged (29). The extensive use of RDV has been restricted by the necessity of delivery by intravenous infusion, which requires access to qualified health care staff and facilities. The FDA and EUA recently authorized two oral antivirals, MPV and Paxlovid, available to specific outpatient populations, which are likely to have a positive impact on public health by reducing the duration of the disease and preventing hospitalization of high-risk individuals. However, cost and accessibility will ultimately determine their global utility. We utilized the K18-hACE2 mouse model, as it is more susceptible to SARS-CoV-2 infection. The K18-hACE2 mouse is an ideal model, as the disease-induced is reminiscent of severe COVID-19 patients with severe pathological changes in both the lung and brain, morbidity, and mortality, as previously reported (18, 22). Upon SARS-CoV-2 infection, these mice showed a sensitive and rapid response to viral replication, clinical parameters, weight loss by 4 or 5 DPI, and lethality by 7-8 DPI, whereas other models exhibit a mild response (30). As described in previous studies (31, 32), all NTV, RDV, and MPV-mono-treated animals also exhibited similar dynamics in terms of weight loss, viral titer, pulmonary function, lung pathology, and virus-induced death. Comparable weight loss was observed between mock and RDV-treated animals, whereas NTV and MPV-treated animals had lower body weight loss and higher body weight recovery, with higher significance for MPV. Furthermore, mice treated with MPV had a lower viral burden and sub-genomic viral RNA level than mice treated with other drugs, which was consistent with a previous report and indicated the ability of MPV to reduce virus replication (33). As viral burden and clinical pathology correlate, our study is also reminiscent of this phenomenon and showed that mice treated with MPV exhibited decreased clinical severity, with improvement in the clinical score, pathophysiological changes, and viral distribution compared with those in other treatment groups. According to several studies, NTV and RDV antivirals exhibited survival rates of 36% and 21%, respectively, while MPV-treated animals had approximately 7 and 22% higher survival rates than NTV and RDV-treated animals, respectively animals (6, 26, 34). Recent clinical studies suggest that RDV does not play a significant role in mortality, but it still plays a major role in reducing the duration and severity of illness, an important outcome when hospitals are overcrowded with patients with COVID-19 (35-38). In support of these studies, our *in vivo* study also clearly illuminates the survival benefits unrelated to RDV monotherapy, which resulted in only 21% survival, although it may be of clinical significance because it has yielded improved weight loss recovery and clinical scores. Collectively, we confirmed that the three antivirals have therapeutic significance against SARS-CoV-2 pathogenesis in the K18-hACE2 mouse model, with lower protection afforded by RDV and higher protection afforded by MPV.

The antiviral potency of the drug combinations against SARS-CoV-2 viral infection *in vitro* has been depicted (17, 39); however, there are limited studies available on FDA-approved drugs *in vivo* models. Since the three individual drugs studied showed therapeutic effects, we further evaluated the therapeutic efficacy of the drugs in combination with each other to consider whether the combination could show synergistic antiviral activity compared to that with monotherapy. This study was the first to examine the value of combination therapy with NTV-RDV and NTV-MPV combinations against SARS-CoV-2 infection in an animal model and, more importantly, the ability of combination therapy to extend the treatment window. Furthermore, potential side effects with a high dose and poor therapeutic potency with a low dose must be considered. These limitations can be overcome by the application of the synergistic action of the currently employed suboptimal dose of the NTV-MPV combination. We reported here that the suboptimal dose (20 mg/kg body weight) drug combinations produced the same pattern in protection against weight loss; however, NTV-MPV yielded a significantly better clinical score than NTV-RDV, emphasizing the importance of the NTV-MPV combination in preventing the clinical progression of the disease. Interestingly, the clinical score associated with NTV-RDV was significantly lower than that with NTV or MPV monotherapy, which is a considerable factor when recommending this combination therapy for the treatment of SARS-CoV-2 infection. The favipiravir and MPV combination has recently been reported to reduce the viral load better than monotherapy in SARS-CoV-2-infected mice (40). The reduced viral load with the NTV-MPV combination is superior to that in animals treated with the other drugs at 5 DPI, representing the clinical significance of this combination in the clearance of the virus. Combinations of different drugs, such as ribavirin-favipiravir and oseltamivir-azithromycin in Lassa virus and influenza-infected mice, respectively, substantially extended the survival period and led to a better survival rate compared to those with monotherapy (41, 42). Mice treated with the NTV-MPV combination had a significantly reduced viral distribution in the lungs as well as in the brain and ultimately exhibited 80% survival, while mice treated with other drugs exhibited a lower (<45%) survival rate. Moreover, this achieved survival rate is more pronounced than expected from the additive activity of NTV or MPV monotherapies. Together, these data reveal that the administration of the NTV-MPV combination can considerably diminish virus replication in the early onset of the disease, where the predominant viral load is usually expected, limiting deleterious COVID-19 infectious events.

Although the overall results of this study are encouraging for their clinical significance, some limitations are also notable. Our results support the need for the characterization of the pharmacodynamic and pharmacokinetic properties, mainly the toxicity, of the NTV-MPV combination in appropriate animal models and humans before treating COVID-19 patients. Combination therapy has been associated with the acquisition of fewer genomic mutations; however, there is an advantage to examining the genetic mutation of the SARS-CoV-2 virus against this combination therapy. Our research demonstrated that three studied antiviral drugs had therapeutic potential by diminishing the viral burden, rescuing the lung and brain pathology, and ultimately improving life expectancy by 20-40%. The NTV-MPV combination was highly protective against lethal SARS-CoV-2 infection and yielded a survival rate of approximately 80% by controlling all pathological features associated with morbidity and mortality. Combination therapy with different drugs affects different portions of the viral life cycle and may be a beneficial strategy based on their mechanisms of action against SARS-CoV-2 infection. We achieved such a potential effect while applying a combinatorial approach with NTV-MPV combination therapy and presented *in vivo* evidence of the potential utility of this combination in treating SARS-CoV-2 patients.

## MATERIAL AND METHODS

### Ethics statement

The care and maintenance of the animals and animal housing followed the recommendations and guidelines provided by the Ministry of Food and Drug Safety, Republic of Korea. This study was approved by the institutional animal care and use committee of Chungbuk National University, Republic of Korea (CBNUA-1659-22-01), and all experimental protocols strictly adhered to committee guidelines.

### Cells and virus

Vero E6 cells (African green monkey kidney cells, ATCC®, cat# CRL-1586(tm)) were maintained in Dulbecco’s modified Eagle’s medium (DMEM) containing 10% (w/v) fetal bovine serum (FBS), 4.0 mM L-glutamine, 110 mg/L sodium pyruvate, 4.5 g/L D-glucose and 1% antibiotic-antimycotic at 37 °C and 5% CO2.

The virus Beta-CoV/Korea/KCDC03/2020 (NCCP43326) used for this study was obtained from the National Culture Collection for Pathogens, Republic of Korea. It was propagated in Vero E6 cells maintained in DMEM supplemented with 2% FBS and 1% antibiotic-antimycotic at 37 °C and 5% CO2. All the infection work related to SARS-CoV-2 was carried out in the biosafety level 3 (BSL3) and animal BSL3 (ABSL3) facilities at Chungbuk National University per the institutional ethical committee standards.

### Compound sources

Nirmatrelvir and molnupiravir were purchased from MedChemExpress (Monmouth Junction, USA). Remdesivir was purchased from Ambeed (Illinois, USA). Stock solutions of all these antiviral drugs were prepared in 2% DMSO and 20% sulfobutylether-β-cyclodextrin (MedChemExpress) in 0.9% saline and stored at -20 °C.

### SARS-CoV-2 infection in K18-hACE2 mice and treatment regimen

Eight-week-old female C57BL/6 Cg-Tg(K18-ACE2)2Prlmn/J mice were obtained from Jackson Laboratory, USA, and maintained in a pathogen-free ABSL3 facility at the desired humidity, temperature, and day/night conditions. All experimental mice were divided into 7 groups with 14 mice in each group: group 1: mock-treated, group 2: NTV (20 mg/kg), group 3: RDV (20 mg/kg), group 4: MPV (20 mg/kg), group 5: NTV (20 mg/kg) + RDV (20 mg/kg), group 6: NTV (20 mg/kg) + MPV (20 mg/kg), and group 7: vehicle. Mice were anesthetized by isoflurane inhalation and inoculated with 5 MLD50 of Beta-CoV/Korea/KCDC03/2020 in 50 µL. NTV and MPV were administered orally for treatment, and RDV was given intraperitoneally twice daily (b.i.d.) for five days starting at 6 hours post-infection (dpi). The combinatorial treatment of MPV or RDV was applied after administering NTV.

### Weight, clinical score, and survival

The body weight changes and survival of SARS-CoV-2-infected mice were monitored daily for 14 days. The mice were also monitored and scored for clinical symptoms daily up to 14 days post-infection (DPI) via a clinical scoring system used to monitor disease progression and establish human endpoints (18). The categories included body weight, appearance (fur, eye closure), activity, and movement, which were evaluated according to standard guidelines with a maximum score of 11. The mice were sacrificed when they lost ≥25% of their initial pre-infection body weight. Lung and brain tissues were collected at 5 DPI (n = 5/group) to determine viral titers, viral RNA copies, histopathology (hematoxylin and eosin; H&E), and immunohistochemistry and stored at -80 °C for further use.

### Infectious viral titer estimation

The coinfection method was used to estimate the viral titer in homogenized lung and brain tissues. Briefly, the tissues were homogenized with bead disruption and centrifuged at 12,000 rpm for 5 min. The 10-fold serially diluted virus-containing samples were then used to infect 0.2 × 10^6^ Vero E6 cells per well in a 96-well plate and cultured in a 5% CO2 incubator at 37 °C for 4 days or until an apparent cytopathic effect (CPE) was detected and further visualized by 0.2% crystal violet staining. The titer was calculated according to the Reed and Muench method and was expressed in log10 50% tissue culture infection dose (TCID50) per mg of tissue (43).

### Quantitation of SARS-CoV-2 RNA

Isolation of RNA from homogenized lung and brain tissue was performed with a QIAamp Viral RNA Kit (QIAGEN) per the manufacturer’s instructions, with slight modifications. Complementary DNA conversion and RT–qPCR were accomplished by a Bio-Rad Real-Time PCR System using a VBScript II One-Step RT–qPCR Probe Kit (Seegene Inc.). Viral genome copies of each of the RdRp and N genes of the SARS-CoV-2 viral RNA were counted on the Ct values using the matrix gene-based real-time reverse transcriptase-polymerase chain reaction (44).

### Histopathology and immunohistochemistry analysis

Mouse lung and brain tissues collected at 5 DPI were fixed with 10% formalin neutralization buffer (Sigma Aldrich). The paraffin-embedded tissue sections were stained with H&E for evaluation of the histopathological score by a pathologist. The semiquantitative scoring system (0 - normal, 1 - mild, 2 - moderate, and 3 - severe) was applied by examining the tissues under digital microscopy as described previously (18, 45, 46). The cumulative scoring system was applied by considering perivascular inflammation, bronchiolar epithelial necrosis, bronchiolar epithelial necrosis regeneration, bronchiolar inflammation, alveolar inflammation, and perivascular edema. Immunoreactivity in the lung tissues was assessed using a rabbit anti-SARS-CoV-2 Nucleocapsid pAb as the primary antibody (Sinobiological, China) and further processed using a Ventana Discovery Ultra (Roche, USA) system.

### Statistical analysis

All statistical data were analyzed using GraphPad Prism 9 (GraphPad, San Diego, USA) and SPSS 26 (IBM SPSS, New York, USA) software. Statistical significance among different groups was determined using a one-way analysis of variance (ANOVA) followed by Tukey’s multiple comparisons test. Survival analysis was performed using the Kaplan–Meier method, and differences were calculated using the log-rank Mantel-Cox test. P values of *p < 0.05, **p < 0.01, and ***p < 0.001 were considered to indicate a significant difference.

## ACKNOWLEDGMENT

This work was supported by grants from the Korea National Institute of Health, the Korea Disease Control and Prevention Agency (4861-312-320-01 to M-. S.S.), and the National Research Foundation of Korea (NRF-2021R1A2C2006961 to M-. S.S. and NRF-2020R1A5A2017476 to M-. S.S.).

## Supplementary figure legends

**Figure S1:**

Lethal infection dose optimization of SARS-CoV-2 virus-infected K18-hACE2 transgenic mice. **(A)** The progression of body weight loss was measured daily until 14 DPI in mice infected with different doses of the Beta-CoV/Korea/KCDC03/2020 virus. Weight loss was measured as a percentage of the initial body weight at day 0. **(B)** Kaplan–Meier survival plot of mice inoculated with different doses of the Beta-CoV/Korea/KCDC03/2020 virus.

**Figure S2:**

Dose-optimization studies for low-dose therapeutic efficacy determination using hACE-2 transgenic mice infected with the Beta-CoV/Korea/KCDC03/2020 virus. **(A)** Depicted percentage of the body weight measured at 14 DPI in the infected mice therapeutically treated with 10 mg/kg and 20 mg/kg body weight drugs. **(B)** Kaplan–Meier survival plot of the mice treated with 10 mg/kg and 20 mg/kg body weight drugs.

**Figure S3:**

The weight comparison of the groups treated with mono and combination drugs at 7 DPI. A significant difference was detected between the nirmatrelvir-molnupiravir and mock-treated group (One-way ANOVA with Turkey’s multiple comparison tests).

**Figure S4:**

Histopathology and immunochemistry analysis of SARS-CoV-2-infected mice treated with 20 mg/kg body weight different drugs. **(A)** Pathological changes and virus distribution were measured at 14 DPI, and images are shown at both low (5X) and high (20X) power resolution. **(B)** Neuronal death and virus distribution were measured at 14 DPI, and images are shown at both low (10X) and high (20X) power resolution.

